# Dissociable Neural Responses to Conditioned Social Threats are Modulated by Spatial Proximity in PTSD and MDD

**DOI:** 10.64898/2026.07.10.737814

**Authors:** Nick Steele, Leonel Rangel-Jimenez, Dash A. Watts, LeeAnne Tunstall, Jenna A. Beakas, Ahmed Hussain, Ashley Huggins, Delin Sun, Kevin S. LaBar, Rajendra A. Morey

## Abstract

The brain’s response to potential threats is shaped by the egocentric spatiotemporal distance to the threat. While distal threats recruit evaluative, cognitive-fear networks for strategic planning, proximal threats engage evolutionarily conserved reactive-fear circuitry to mount immediate defensive responses. Understanding this dual neural architecture is clinically relevant, as the spatial proximity to a traumatic event predicts subsequent psychiatric symptom severity and recovery. Despite the well-characterized disruptions to threat circuitry in posttraumatic stress disorder (PTSD) and major depressive disorder (MDD), how these networks respond to threats at distinct spatial distances has yet to be investigated in these disorders. Utilizing 3D virtual reality technology, we implemented a spatially-modulated fear conditioning paradigm by presenting human avatars at proximal (peripersonal) and distal (extrapersonal) distances, paired with shock, to trauma-exposed participants (*n* = 50) during functional MRI (fMRI). We modeled associations between PTSD and MDD symptom severity and task-evoked hemodynamic responses within cognitive-fear and reactive-fear threat networks. Our results support a transdiagnostic disruption of thalamic responses to proximal threats, driven by heightened activation to the safety stimulus and stronger deactivation to the threat stimulus. MDD showed a disorder-specific effect of disrupted activity across cognitive-fear and social cognition regions in response to proximal social threats, and illness severity-by-proximity effects across motor regions. PTSD symptom severity was uniquely associated with hyperactivity of the amygdala to distal threats. Transdiagnostic disruption of connectivity between the dorsal precuneus and several subcortical structures followed a disorder-specific gradient across the anterior-posterior axis. While thalamic disruptions to threats represent a shared transdiagnostic effect, PTSD and MDD are distinguished by differences in amygdala hyperactivation and cognitive-fear network hypoactivation, respectively.

## Introduction

Adaptive responding to environmental threats is critical for survival. To facilitate the most effective and appropriate response to ecological threats, the brain is thought to employ specialized threat processing networks that are differentially recruited based on the spatiotemporal gradient of threat proximity. The distinction between proximal and distal threat processing has been best characterized by the predatory imminence model, originating from behavioral ecology, which proposes that organisms shift from deliberative, goal-directed cognition to automatic, reflexive defensive behaviors as a predator approaches (1). So-called cognitive-fear circuits respond to more distant threats and consist of evaluative processing regions of the brain. This system includes the ventromedial prefrontal cortex (vmPFC), dorsolateral PFC (dlPFC), posterior cingulate cortex (PCC), precuneus, inferior parietal lobe, amygdala, and hippocampus (2,3). Conversely, reactive-fear circuits are evolutionarily conserved and respond to threats that are spatially and temporally near, including the midbrain and periaqueductal gray (PAG), thalamus, anterior cingulate cortex (ACC), and anterior insula. Meta-analysis of traditional fear conditioning studies demonstrate that reactive-fear regions show greater activation to the conditioned stimulus (CS+), while cognitive-fear regions display greater activation in response to the safety stimulus (CS-) (4).

While threat processes have been well studied in the human brain, only a small number of neuroimaging studies have modeled threat across spatiotemporal scales. Most investigations modeling threats across space and time have used chase-and-capture video games in which healthy participants navigate through a virtual environment by controlling a simple shape-avatar representing the participant is approached by a predator, which is represented by a different geometric shape (2,3). Only recently have more naturalistic threat stimuli been incorporated at distinct spatial distances. Faul et al. used MRI-compatible 3D technology to examine neural responses to human avatars paired with shock presented at near and far distances to assess spatially-modulated threat acquisition and extinction (5). Their findings further support the distinction between cognitive-fear and reactive-fear processing regions.

Understanding these processes is relevant for trauma-based clinical disorders, as the spatial proximity of the trauma survivor to the threat predicts the likelihood of developing post-trauma psychopathology and subsequent clinical symptom severity (6,7). Post-trauma behavioral changes most commonly present as posttraumatic stress disorder (PTSD) and major depressive disorder (MDD). PTSD symptoms include hypervigilance to potential threats, avoidance of trauma-related threat cues, negatively altered mood and cognitions, and the persistent re-experiencing of trauma memories (8). Associations between elevated PTSD symptoms and altered threat responding have been well established, but how distinct spatiotemporal threat circuits are differentially impacted by PTSD symptoms is not well understood. Evidence from resting-state connectivity studies implicates spatiotemporal threat circuitry, such as the PAG, in PTSD pathophysiology (9,10). Elevated amygdala reactivity in response to threatening stimuli has also been consistently reported in PTSD (11,12). Yet, whether amygdala hyperreactivity is elicited by both near and far threats, or is preferential to one over the other, has not been explored. MDD is highly comorbid with PTSD and is characterized by robust attentional biases towards negatively valenced and threat-related stimuli (13,14). While PTSD and MDD symptoms display both overlapping and unique patterns of disrupted connectivity within threat circuitry (15), neural responses in MDD during threat conditioning remain under-investigated.

We evaluated associations between PTSD and MDD symptom severity and threat processing of *near* and *far* social threats presented during *early* and *late* acquisition. Trauma-exposed participants completed a function MRI (fMRI) fear conditioning paradigm in which human characters were presented via 3D virtual reality at peripersonal and extrapersonal distances, with and without paired shocks. Hemodynamic responses within cognitive-fear and reactive-fear regions were examined as a function of symptom severity during both acquisition phases. We hypothesized that higher disorder severity would disrupt responses in both circuits. The amygdala exhibits hyperactivation in PTSD and has been implicated in the exaggerated threat reactivity and fear generalization characteristic of the disorder (11,12); we therefore predicted that greater PTSD symptom severity would be associated with heightened amygdala responses, either to proximal or distal threats. Conversely, the dlPFC supports the cognitive appraisal and top-down regulatory processes engaged by distal threats, and is reliably disrupted in MDD (16,17). We therefore predicted that greater MDD symptom severity would be associated with reduced dlPFC engagement during distal threats.

## Methods

### Sample

Fifty trauma-exposed participants were recruited through the Durham Veterans Affairs Health Care System (VAHCS) and compensated for their participation at $25 per hour. Recruitment was limited to post-2001 military veterans. Participants were excluded for any neurological, developmental, or psychiatric comorbidities (excluding PTSD and MDD), a history of head trauma resulting in loss of consciousness or hospitalization, or MRI contraindications. To minimize the influence of confounds, recruitment was capped such that current psychotropic medication use did not exceed 25% of the total sample. All participants provided written informed consent, and all study procedures were approved by the VAHCS and Duke University Health System Institutional Review Boards. Demographic and clinical data are presented in *Table 1*. Clinical variable correlations are reported in *Figure S1*.

**Table 1.** Demographic and Clinical Variables.

*Demographic and Clinical Variables*
| **Variable** | ***N* = 50** |
| --- | --- |
| PTSD Diagnosis (CAPS-5) | *n* = 14 |
| PTSD Severity (CAPS-5) | 16.5 (14.8) |
| Re-experiencing symptoms | 3.8 (4.0) |
| Avoidance symptoms | 2.2 (2.2) |
| Negative Mood/Cog symptoms | 5.4 (5.9) |
| Hyperarousal symptoms | 5.1 (4.4) |
| MDD Diagnosis (BDI-II) | *n* = 28 |
| MDD Severity (BDI-II) | 15.9 (11.7) |
| Cognitive symptoms | 5.0 (4.7) |
| Somato-Affective symptoms | 10.9 (7.7) |
| Age | 45.2 (8.7) |
| Sex |  |
| Female | 17 (34%) |
| Male | 33 (66%) |

Mean (SD); *n* (%)

### Procedure and Materials

The study was conducted across two sessions approximately 24 hours apart. On Day 1, participants underwent habituation, acquisition, and extinction runs. On Day 2, participants underwent one extinction-recall run. The current analyses focus exclusively on the Day 1 acquisition to evaluate threat acquisition processes. Conditioned stimuli (CS) and unconditioned stimuli (US) consisted of four distinct male human avatars displayed at peripersonal (1.0 m in virtual space) or extrapersonal (5.0 m in virtual space) distances while participants passively moved forward down an alleyway (0.3 m/s) and paused during each 6 s avatar presentation. Stimuli included one near stimulus paired with shock (CS+_Near_), one far stimulus paired with shock (CS+_Far_), one near stimulus not paired with shock (CS-_Near_), and one far stimulus not paired with shock (CS-_Far_). Each of the stimuli were presented 10 times, resulting in 40 total trials. Visual stimuli were initially presented using a MagnetVision 3D television (*n* = 39) and were later upgraded to Nordic Neuro VisualSystem HD 3D goggles (*n* = 11) during data collection. All participants had normal or corrected-to-normal vision. Electrical stimulation was paired with the CS+_Near_ and CS+_Far_ stimuli on 60% of trials such that the avatar presentation was co-terminated with electrical stimulation. Stimulation intensity was individually calibrated via a step-up procedure in which participants directed voltage increases until they considered the stimulations to be “highly annoying but not painful”. Electrical stimulation was applied to the right wrist through a MP-150 BIOPAC system (BIOPAC Systems) using the STM200 Constant Voltage Stimulator.

Immediately following MRI scanning, participants provided behavioral ratings evaluating subjective danger, arousal, and valence of each avatar presented during the fear conditioning paradigm (*Figure S2*). Psychiatric symptom severity was assessed in accordance with the Diagnostic and Statistical Manual of Mental Disorders-5 (DSM-5) (8) criteria. MDD symptomatology was assessed using the Beck Depression Inventory II (BDI-II) (18) and PTSD symptomatology with the Clinician-Administered PTSD Scale for DSM-5 (CAPS-5) (19).

### US Expectancy Ratings

Using an MRI-compatible button box, participants rated each character presentation using their left hand on the perceived likelihood of receiving a shock (shock expectancy) on a scale from 1 to 4, where 1 indicates “definitely unlikely” (little finger), 2 “somewhat unlikely” (ring finger), 3 “somewhat likely” (middle finger), and 4 “definitely likely” (index finger). These ratings served as a subjective behavioral measure of fear learning. Technical issues with the button box resulted in missing ratings for 16 participants, resulting in a sample size of *n* = 34 for these analyses.

### Task-Modulated Functional Connectivity

Seed-based task-modulated functional connectivity was assessed using generalized psychophysiological interaction (gPPI) analysis. Seed timeseries were deconvolved to estimate the underlying neural signal, then multiplied by the task regressor for each condition to generate PPI terms. The deconvolved physiological timeseries, PPI terms for each task condition, 6 motion parameters, and 10 aCompCor components were entered as regressors in a seed-to-whole-brain voxelwise GLM to estimate task-modulated functional connectivity.

Five bilateral seeds were selected based on their established roles in fear conditioning and post-trauma symptomatology: the amygdala, anterior hippocampus, posterior hippocampus, PAG, and vmPFC (2,3,5). The amygdala and hippocampus were defined using the Automated Anatomical Labeling (AAL) atlas, with the hippocampus divided into anterior and posterior portions at y = −24 mm in MNI space. The PAG was derived from the Harvard-Oxford probabilistic brainstem parcellation (thresholded at ≥ 30%) and further restricted to z > ™24 mm and y < −22 mm in MNI space (5). The vmPFC was defined as the union of the subcallosal cortex and frontal medial cortex from the Harvard-Oxford probabilistic atlas, thresholded at ≥ 50%.

### Statistical Analysis

Behavioral data, including shock expectancy ratings and post-scan ratings of stimulus danger, arousal, and valence, was assessed using an analysis of variance (ANOVA). A repeated-measures 2 × 2 factorial design was implemented to test for associations with stimulus threat (CS+ vs. CS-) and proximity (near vs. far).

Brain data were analyzed by fitting first-level beta maps with linear mixed-effects models using AFNI’s 3dLMEr. We implemented two analytical frameworks. The first examined predefined threat contrasts during both the early acquisition (first five trials) and late acquisition (last five trials) test phases, yielding four specific contrasts: CS+_Near_ > CS-_Near_ and CS+_Far_ > CS-_Far_ for early and late trials. This framework was used to test both activation and gPPI connectivity data. For these analyses, the first-level contrast beta map served as the dependent variable. Symptom severity, age, and sex were entered as fixed effects, with MRI scanner and visual display equipment included as random intercepts. The model was run separately using PTSD and MDD symptom severity. Results using a model including both PTSD and MDD severity is reported in *Table S3*. Results for PTSD and MDD diagnosis are reported in *Figure S3* and *Figure S4*, respectively.

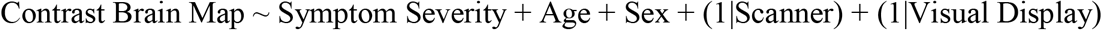

The second framework evaluated neural activation using the first-level beta maps of the individual conditions (CS+_Near_, CS-_Near_, CS+_Far_, CS-_Far_) by modeling interactions between symptom severity, threat, and proximity. We examined the three-way interaction (Symptom Severity × Threat × Proximity) as well as the constituent two-way interactions. To account for multiple trial types per subject, a subject-level random intercept was added. These interaction models were run separately for the early and late acquisition phases:

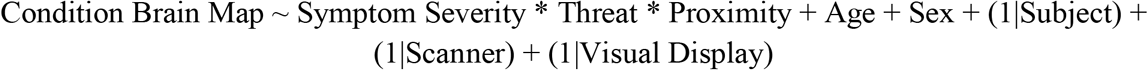

A gray matter mask consisting of near and far threat processing regions was applied to the results. Hypothesized regions from the threat imminence literature included the cerebellum, amygdala, hippocampus, PAG, thalamus, cingulate cortex, vmPFC, dlPFC, precuneus, precentral gyrus, premotor cortex, anterior insula, posterior supramarginal gyrus (pSMG), and angular gyrus (AG) (2,3,5). Results within the thalamus were compared with the Morel atlas (20,21) to determine which nuclei overlapped with the significant voxels.

## Results

### Shock Expectancy and Post-Scan Stimulus Ratings Indicate Successful Threat Learning

Shock expectancy ratings indicated successful learning of threat contingencies (*Figure 2A*). Overall, the CS+ stimuli elicited significantly higher shock expectancy compared to the CS-stimuli (*F*(1,33) = 13.34, *p* < 0.001), and this successful discrimination did not significantly interact with spatial distance (*F*(1,33) = 2.824, *p* = 0.102).

**Figure 1.**
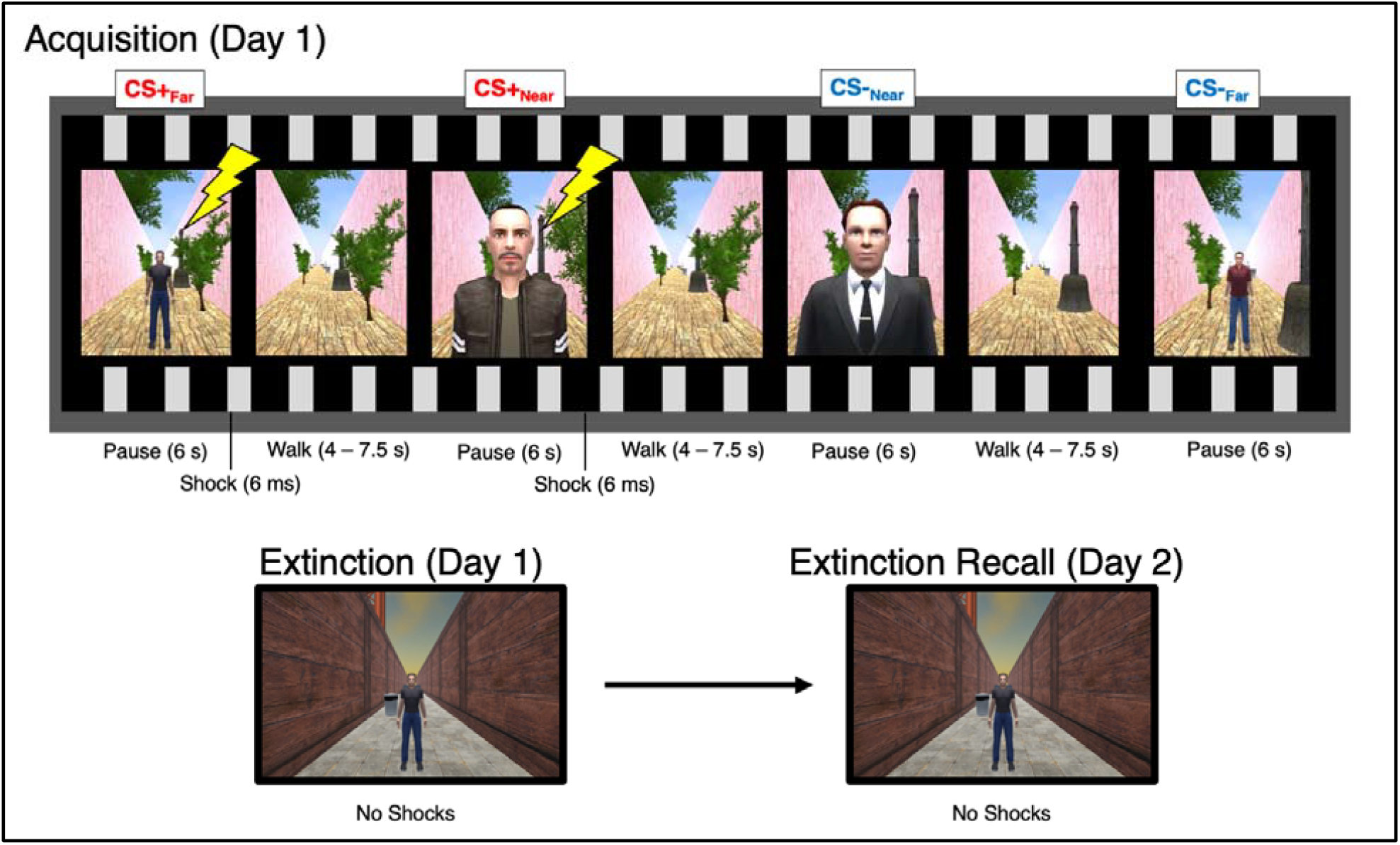
Experimental design. An example of the trial sequence for the acquisition virtual reality run is shown. Similar trial sequences were used for the extinction and extinction-recall runs, but a different context (background) was used and no shocks were used. In the present study, only data from the acquisition phase is reported.

**Figure 2:**
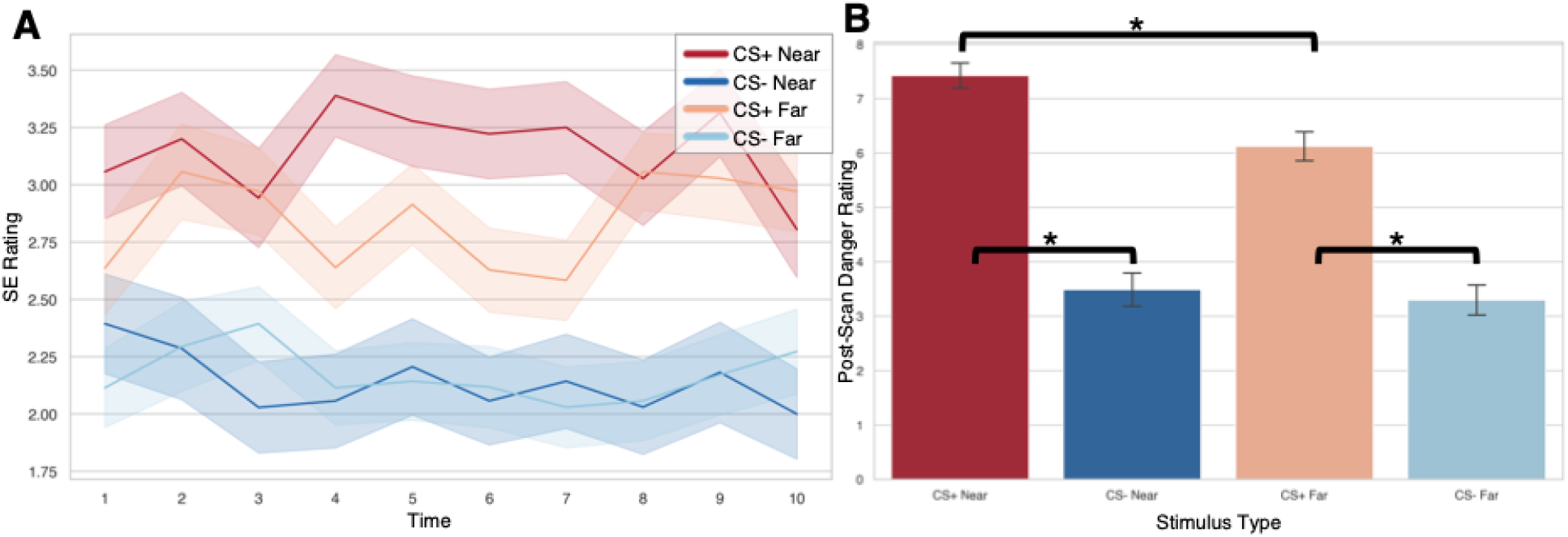
Behavioral data results. **(A)** Shock expectancy ratings (*n* = 34). **(B)** Post-scan stimulus danger ratings (*n* = 50). For each avatar, participants were asked “Is this person safe or dangerous?” and responded on a 9 point Likert scale with responses ranging from “Very Safe” to “Very Dangerous”. Both shock expectancy and danger ratings showed significant effects of threat, with significantly greater ratings for the CS+ compared to the CS-stimuli, suggesting successful learning of threat contingencies. * = *p* < 0.05.

Post-scan stimulus danger ratings revealed significant main effects of threat (*F*(1, 46) = 114.76, *p* < 0.001) and proximity (*F*(1, 46) = 9.48, *p* = 0.003) as well as a significant threat-by-proximity interaction (*F*(1, 46) = 5.21, *p* = 0.027). (*Figure 2B)*. Post-hoc t-tests were performed and showed that participants rated the CS+_Near_ (*M* = 7.43, *SD* = 1.58) and CS+_Far_ (*M* = 6.13, *SD* = 1.83) as significantly more dangerous than the CS-_Near_ (*M* = 3.49, *SD* = 2.10, *t* = 9.28, *p* < 0.001) and CS-_Far_ (*M* = 3.30, *SD* = 1.89, *t* = 7.64, *p* < 0.001), respectively. The CS+_Near_ was rated as significantly more dangerous than the CS+_Far_ (*t* = 4.71, *p* < 0.001), whereas no significant difference was observed between the CS-_Near_ and CS-_Far_ (*t* = 0.48, *p* = 0.63). The same patterns were observed for both arousal (*Table S1*) and valence (*Table S2*) ratings.

Linear regression models adjusting for age and sex revealed no robust associations between PTSD or MDD severity and threat discrimination indices for either shock expectancy or post-scan evaluations. A nominal positive correlation between PTSD severity and CS+_Near_ arousal ratings (*r* = 0.35, *p* = 0.031) did not survive covariate adjustment (*p* = 0.18).

### PTSD Severity Exhibits Distinct Associations with Near and Far Threats

PTSD symptom severity was tested for associations with neural threat processing by examining CS+ > CS-contrasts during early and late acquisition phases. During late acquisition, higher PTSD severity predicted reduced thalamic activity in the ventral posterolateral (VPL) nucleus extending into the anterior pulvinar (PuA) in response to near threats. Additionally, greater PTSD severity predicted higher amygdala activity, extending into the anterior hippocampus, in response to far threats (*Table 2*).

**Table 2.** Univariate Activation Results: PTSD and MDD Severity.

| **PTSD Severity** |  |  |  |  |  |  |
| --- | --- | --- | --- | --- | --- | --- |
| **Region** | **Distance** | **Acquisition Phase** | **Volume (mm<sup>3</sup>)** | **Mean Z** | **Max Z** | **Peak MNI (x, y, z)** |
| VPL Thalamus L.* | Near | Late | 272 | -3.73 | -4.67 | -16, -22, 0 |
| Amygdala R.* | Far | Late | 240 | 3.90 | 4.66 | 30, -10, -20 |
| **PTSD Severity-by-Sex** |  |  |  |  |  |  |
| **Region** | **Distance** | **Acquisition Phase** | **Volume (mm<sup>3</sup>)** | **Mean Z** | **Max Z** | **Peak MNI (x, y, z)** |
| vmPFC | Far | Early | 728 | 3.91 | 4.64 | 0, 16, -20 |
| Cerebellum Crus I L. | Far | Early | 544 | 3.75 | 5.04 | -32, -78, -30 |
| **PTSD Severity-by-Threat** |  |  |  |  |  |  |
| **Region** | **Distance** | **Acquisition Phase** | **Volume (mm<sup>3</sup>)** | **Mean Z** | **Max Z** | **Peak MNI (x, y, z)** |
| Mediodorsal Thalamus* | – | Early | 152 | -3.78 | -4.72 | 2, -12, 4 |
| Angular gyrus R. | – | Late | 376 | -3.89 | -4.92 | 56, -54, 34 |
| PuA Thalamus L.* | – | Late | 301 | -3.61 | -4.22 | -12, -24, 2 |
| Centromedial Thalamus R.* | – | Late | 160 | -3.60 | -4.12 | 12, -18, 2 |
| Mediodorsal Thalamus* | – | Late | 160 | -3.67 | -4.02 | 0, -12, 4 |
| **PTSD Severity-by-Proximity** |  |  |  |  |  |  |
| **Region** | **Distance** | **Acquisition Phase** | **Volume (mm<sup>3</sup>)** | **Mean Z** | **Max Z** | **Peak MNI (x, y, z)** |
| dIPFC R. | – | Late | 312 | -3.68 | -4.38 | 32, 22, 42 |
| **MDD Severity** |  |  |  |  |  |  |
| **Region** | **Distance** | **Acquisition Phase** | **Volume (mm<sup>3</sup>)** | **Mean Z** | **Max Z** | **Peak MNI (x, y, z)** |
| Midbrain* | Near | Early | 160 | -3.82 | -4.74 | 6, -32, -22 |
| VPL Thalamus L.* | Near | Late | 200 | -3.61 | -4.20 | -18, -20, 8 |
| dIPFC R. | Near | Late | 1176 | -3.91 | -5.10 | 38, 24, 36 |
| Angular gyrus R. | Near | Late | 816 | -3.62 | -4.70 | 40, -56, 46 |
| PCC | Near | Late | 408 | -3.65 | -4.66 | 0, -38, 36 |
| Premotor L. | Far | Late | 344 | -3.72 | -4.64 | -50, 10, 36 |
| **MDD Severity-by-Sex** |  |  |  |  |  |  |
| **Region** | **Distance** | **Acquisition Phase** | **Volume (mm<sup>3</sup>)** | **Mean Z** | **Max Z** | **Peak MNI (x, y, z)** |
| Cerebellum Crus II R. | Near | Early | 784 | -4.19 | -6.58 | 6, -82, -38 |
| Precuneus | Near | Early | 376 | -3.55 | -4.22 | -2, -80, 42 |
| Motor cortex R. | Near | Early | 464 | -3.92 | -4.93 | 6, -26, 80 |
| vmPFC. | Far | Early | 280 | 3.56 | 4.20 | -4, 14, -8 |
| Motor cortex R. | Far | Late | 352 | 3.77 | 4.53 | 36, -12, 58 |
| **MDD Severity-by-Threat** |  |  |  |  |  |  |
| **Region** | **Distance** | **Acquisition Phase** | **Volume (mm<sup>3</sup>)** | **Mean Z** | **Max Z** | **Peak MNI (x, y, z)** |
| Precuneus | – | Late | 568 | -3.91 | -4.95 | -2, -60, 60 |
| dIPFC R. | – | Late | 536 | -3.90 | -5.07 | 40, 24, 34 |
| pSMG R. | – | Late | 312 | -3.69 | -4.39 | 48, -38, 48 |
| PuM Thalamus R.* | – | Late | 200 | -3.79 | -4.66 | 12, -28, 12 |

MDD Severity-by-Proximity
| Region | Distance | Acquisition Phase | Volume (mm<sup>3</sup>) | Mean Z | Max Z | Peak MNI (x, y, z) |
| --- | --- | --- | --- | --- | --- | --- |
| Cerebellum VIIIB R. | – | Early | 424 | 3.66 | 4.34 | 22, -54, -50 |
| Motor cortex R. | – | Late | 304 | -4.07 | -5.06 | 2, -26, 76 |

Notes: CS+ > CS- contrasts were examined for near and far threats during early and late threat acquisition trials. Positive Z-scores for the sex interactions indicate that the association between symptom severity and BOLD response is significantly more positive for males compared to females ( $\beta_{\text{Male}} > \beta_{\text{Female}}$ ). Negative Z-scores indicate the reverse ( $\beta_{\text{Female}} > \beta_{\text{Male}}$ ). Dashes (–) in the distance column indicate that the respective effect collapsed across both near and far threat conditions. Abbreviations: dlPFC = dorsolateral prefrontal cortex; PCC = posterior cingulate cortex; pSMG = posterior supramarginal gyrus; PuA = anterior pulvinar; PuM = medial pulvinar; vmPFC = ventromedial prefrontal cortex; VPL = ventral posterolateral. \*Regions were small volume corrected (SVC).

Dominance analyses identified which symptom clusters drove these associations. For thalamic activity during late acquisition, the full model explained 43.2% of variance, with avoidance dominating fit (39.6% of R^2^) and negative mood/cognition (23.1%), re-experiencing (19.2%), and hyperarousal (18.0%) contributing moderately (*Figure 3A*); complete dominance analysis confirmed avoidance outperformed all clusters across every submodel. The amygdala model showed similarly high fit (R^2^ = 0.47) but a distinct profile: hyperarousal (32.2%) and negative mood/cognition (31.5%) were primary drivers, with re-experiencing (18.7%) and avoidance (17.7%) moderate (*Figure 3B*); both exhibited conditional and complete dominance over remaining clusters, with substantial incremental R^2^ after accounting for shared variance.

**Figure 3:**
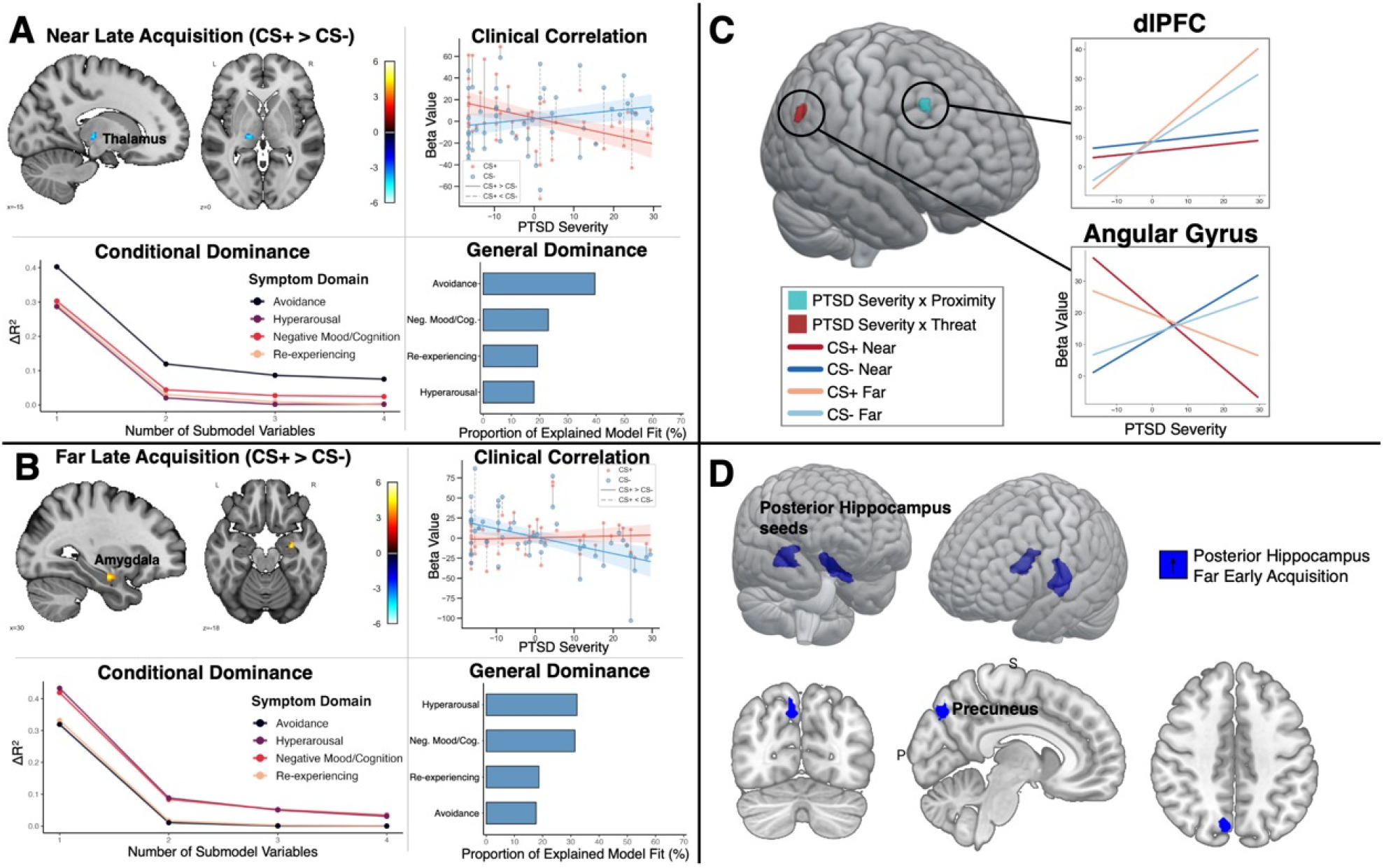
Significant brain associations with PTSD symptom severity. **A)** Higher PTSD severity correlated with reduced thalamic activity in response to near threat during late acquisition. Dominance analysis showed this effect to be predominantly driven by avoidance symptoms. **B)** Higher PTSD severity correlated with increased amygdala activity in response to far threat during late acquisition. Dominance analysis showed this effect to be predominantly driven by hyperarousal symptoms. **C)** Significant interaction effects for PTSD severity-by-proximity and PTSD severity-by-threat interactions (subcortical results not shown). **D)** Task-modulated functional connectivity results. During early acquisition, higher PTSD symptom severity was associated with weaker posterior hippocampus– precuneus connectivity for far threats.

A PTSD severity-by-sex interaction emerged in several regions during early acquisition for far threats, such that more severe PTSD symptoms positively correlated with neural activation among males but negatively correlated among females. Regions included the left cerebellum crus I and the vmPFC. Statistical results are shown in *Table 2*.

PTSD severity-by-proximity interactions during early acquisition emerged in the right dlPFC, with far stimuli driving hyperactivity relative to near stimuli at higher symptom levels (*Figure 3C*). During late trials, a PTSD severity-by-threat interaction appeared in the right angular gyrus, which was hyperactive to CS-relative to CS+ at higher symptom levels. Several thalamic clusters also showed severity-by-threat interactions: within the mediodorsal (MD) nucleus during early trials, and during late trials across three clusters – the medial MD nucleus, right lateral MD/centromedian (CM) nuclei, and left PuA/CM nuclei.

Analysis of gPPI data revealed altered connectivity in response to far threat (CS+_Far_ > CS-_Far_) during early acquisition. PTSD severity correlated with stronger connectivity between the posterior hippocampus and the posterior-dorsal precuneus (volume = 896 mm^3^, max-Z = 4.45, *p*_*bonferroni*_ < 0.05, MNI-peak = [-8, 76, 42]). Dominance analysis showed that variance of connectivity strength (R^2^ = 0.34) was driven predominantly by hyperarousal symptoms (50% of R^2^) (*Table 3*).

**Table 3.** Dominance Analysis Results for gPPI Functional Connectivity: PTSD and MDD Severity.

| PTSD Severity |  |  |  |  |  |  |  |  |  |
| --- | --- | --- | --- | --- | --- | --- | --- | --- | --- |
| Seed Region | Target Region | Distance | Time | R<sup>2</sup> | Adj. R<sup>2</sup> | Re-experiencing | Avoidance | Neg. Mood/Cog. | Hyperarousal |
| Posterior Hippocampus | Precuneus | Far | Early Acquisition | .34 | .25 | .06 (17.1%) | .06 (17.0%) | .05 (14.9%) | .17 (50.0%) |
| MDD Severity |  |  |  |  |  |  |  |  |  |
| Seed Region | Target Region | Distance | Time | R<sup>2</sup> | Adj. R<sup>2</sup> | Cognitive | Somato-Affective |  |  |
| PAG | Precuneus | Near | Early Acquisition | .51 | .47 | .24 (47.7%) | .27 (52.3%) |  |  |
| Amygdala | Precuneus | Far | Early Acquisition | .29 | .23 | .17 (56.1%) | .13 (43.9%) |  |  |
| Amygdala | Precuneus | Far | Early Acquisition | .32 | .26 | .19 (57.9%) | .14 (42.1%) |  |  |

**Note.** R<sup>2</sup> = full model R<sup>2</sup>; Adj. R<sup>2</sup> = adjusted R<sup>2</sup>. Symptom cluster columns display raw dominance R<sup>2</sup> values with the proportion of total model R<sup>2</sup> in parentheses. Age and sex were held constant across all submodels. PAG = periaqueductal gray; Neg. Mood/Cog. = negative mood and cognition.

### MDD Severity Exhibits Distinct Associations with Near Versus Far Threats

Higher MDD severity was associated with widespread altered neural responses across cognitive-fear and reactive-fear regions (*Figure 4A*). During early acquisition trials, more severe MDD symptoms were negatively correlated with near threat (CS+ > CS-) in the midbrain, driven by greater CS-activity and reduced CS+ activity. During late trials, the same effect was observed in the thalamus (VPL nucleus), dlPFC, PCC, and angular gyrus for near threats and the premotor cortex extending into primary motor cortex for far threats.

**Figure 4:**
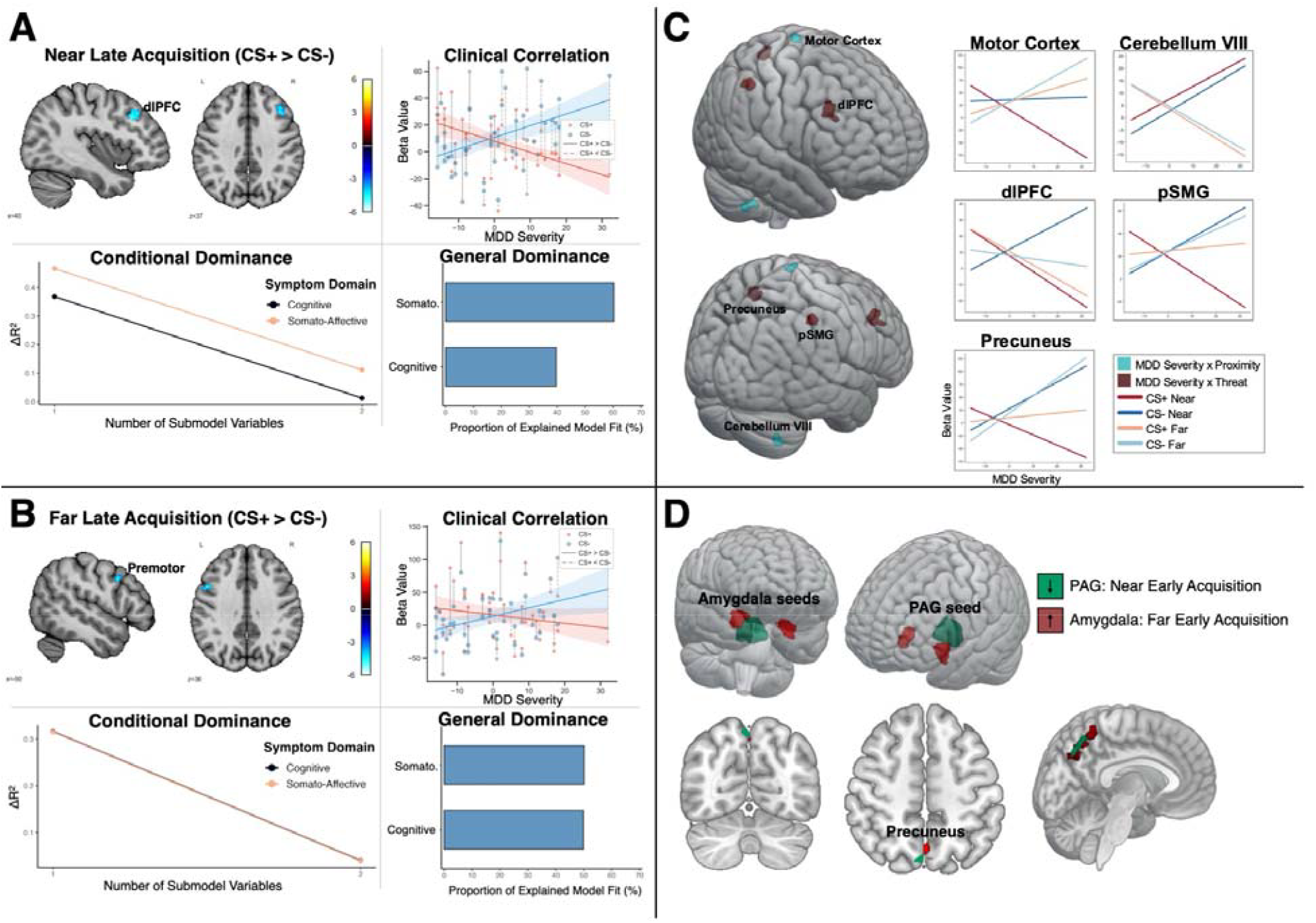
Significant brain associations with MDD symptom severity. **A)** Higher MDD severity correlated with reduced dlPFC activation in response to near threat during late acquisition, and **B)** reduced premotor cortex activation in response to far threats during late acquisition. **C)** Interaction results for MDD severity-by-proximity and MDD severity-by-threat (subcortical results not shown). **D)** Task-modulated functional connectivity (gPPI) results. During early acquisition, higher MDD severity was associated with weaker PAG–precuneus connectivity for near threats, but stronger posterior hippocampus–precuneus connectivity for far threats. Abbreviations: dlPFC = dorsolateral prefrontal cortex, PAG = periaqueductal gray.

Dominance analysis models included BDI cognitive and somato-affective symptom domains with age and sex as covariates. dlPFC and PCC activation was primarily driven by the somato-affective domain, midbrain activity by the cognitive domain, and thalamus, angular gyrus, and premotor regions showed relatively even variance partitioning across both domains.

Several regions showed a MDD severity-by-sex interaction during early acquisition, with positive correlations in females and negative correlations in males. Near threat implicated the right cerebellum crus II, posterior-dorsal precuneus, and dorsal primary motor cortex; far threats implicated the vmPFC. During late acquisition, this sex-dependent pattern reversed in the primary motor cortex for far threats, with males showing a positive and females a negative severity correlation. Results are shown in *Table 2*.

MDD severity-by-proximity interactions emerged in motor regions: right cerebellum VIII (VIIIa/VIIIb) during early acquisition and primary motor cortex during late acquisition. In cerebellum VIII, higher severity correlated with greater near-than far-stimulus activation; the motor cortex showed the opposite pattern, with greater far-than near-stimulus activation. During late trials, right dlPFC, right pSMG, anterior-dorsal precuneus, and medial pulvinar (PuM) clusters showed MDD severity-by-threat interactions, driven by stronger CS™ activation and, particularly for CS+Near, stronger CS+ deactivation (*Figure 4C*).

Finally, MDD symptom severity was associated with disrupted connectivity of the dorsal precuneus during early acquisition. Specifically, higher MDD severity correlated with stronger amygdala–precuneus connectivity for far threats (CS+_Far_ > CS-_Far_) (volume = 600 mm^3^, max-Z = 4.33, *p*_*bonferroni*_ < 0.05) and weaker PAG–precuneus connectivity for near threats (CS+_Near_ > CS-_Near_) (volume = 680 mm^3^, max-Z = -6.26, *p*_*bonferroni*_ < 0.05). Both significant voxel clusters spanned the anterior-posterior axis of the dorsal precuneus (*Figure 4D*). Dominance analysis revealed an even partitioning of variance across the cognitive and somato-affective symptom domains (*Table 3*).

## Discussion

Utilizing a 3D virtual reality environment with fMRI, we show that the spatial proximity of threat reveals distinct pathophysiological correlates for PTSD and MDD symptoms. Behavioral findings indicated successful learning of threat contingencies. Neural findings demonstrated that PTSD symptom severity is associated with reduced thalamic activity in response to proximal threats, while distal threats evoked amygdala hyperreactivity and sex-specific activity alterations among select cognitive-evaluative regions. MDD symptom severity was associated with more widespread disruption across the cognitive-fear network and select reactive-fear regions implicated in mounting defensive responses. These effects were primarily driven by heightened neural activation to non-threat stimuli and reduced activation to threat stimuli with increasing symptom severity.

### Amygdala Hyperreactivity is Specific to PTSD

Amygdala hyperreactivity to conditioned threats in PTSD has been well established (11,12,22). In traditional Pavlovian conditioning models, threat contingency learning is carried out by the basolateral complex of the amygdala (23). Our finding of stronger amygdala activity in PTSD is therefore well supported. Notably, we extend this framework by showing that the pattern of hyperactivity is selectively sensitive to distal threats. The spatial tuning of amygdala hyperreactivity to distal threats aligns with the amygdala’s integration with the cognitive-fear network (2). Furthermore, by partitioning the variance of amygdala activity across PTSD symptom clusters, we showed that this hyperactivity was predominately driven by hyperarousal and negative mood/cognition symptoms. In proposed models of PTSD hyperarousal, heightened amygdala responsivity drives hyperarousal symptoms, which subsequently cascades into affective and cognitive symptoms (24).

### Thalamic Hypoactivity is a Transdiagnostic Effect

The thalamus is a functionally heterogeneous structure whose nuclei relay and modulate sensory information to the cortex, playing a critical role in threat memory encoding and stimulus contingency learning. Several nuclei — MD, CM, VPL, and pulvinar — showed significantly reduced activation with increasing PTSD or MDD severity. In prior work, we reported that PTSD severity is associated with stronger VPL resting-state connectivity with the salience and somatosensory networks, and weaker MD and pulvinar connectivity with sensory and motor cortices (25). Here, thalamic hypoactivation to proximal threats localized to the VPL in both disorders, likely reflecting blunted somatosensory encoding of near threats given the use of shock as the aversive stimulus. Higher-order nuclei (MD, CM, pulvinar) showed heightened responsivity to non-threat stimuli and reduced CS+ activation, with the pulvinar displaying transdiagnostic effects and the MD and CM nuclei specific to PTSD.

### Transdiagnostic and Disorder-Specific Activity Patterns in the Cognitive-Fear Network

Widespread hypoactivity to the CS+ and hyperactivity to the CS-was observed across the cognitive-fear network with increasing MDD symptom severity, including the dlPFC, PCC, and angular gyrus, suggesting increased attunement to safety signals among these regions in MDD. These results align with meta-analysis of fear conditioning studies in healthy samples, which show that cognitive-fear regions display greater activity to the CS-than the CS+ (4). MDD severity interacted with stimulus threat or proximity in the dlPFC, pSMG, and precuneus, and with sex in the vmPFC, cerebellum crus II, and posterior-dorsal precuneus. Prior investigations have highlighted a relationship between trauma exposure and neural responses during safety signal learning in the dlPFC, amygdala, and hippocampus (26). In former military veterans, who form the present cohort, attunement to safety signals is an integral part of patrolling, combat, and enemy engagement. By contrast, most civilian trauma survivors experience trauma as unexpected events not preceded by surveillance for safety.

### Dorsolateral Prefrontal Cortex

Disrupted dlPFC response to threat emerged as a transdiagnostic feature, yet the specific presentation of dysfunction was disorder dependent. Consistent with previous literature detailing dlPFC hypoactivity to negatively valenced stimuli in MDD (16,17,27,28), the dlPFC exhibited a blunted response during late acquisition after threat contingencies were learned. This blunted responsivity to threat occurred alongside hyperactivity to safety. By contrast, PTSD severity interacted with stimulus proximity, which was characterized by dlPFC hyperactivity to distal stimuli, regardless of threat value. This occurred during early acquisition when participants were actively learning threat contingencies. Together, these findings suggest disorder-specific dysfunction in the dlPFC. In PTSD, the dlPFC is hypervigilant to potential threats during threat learning, over-sensitizing the cognitive-fear network to ambiguous distant stimuli. Our findings suggest that in MDD, the dlPFC exhibits a classic blunted response to affect-laden stimuli, but an over-engagement to safety, reflecting enhanced threat evaluation of safe conditions.

### Dorsal Precuneus

The dorsal precuneus exhibited a pattern of disrupted connectivity with several subcortical seeds during early acquisition in both PTSD and MDD following a disorder-specific anterior-posterior gradient, consistent with resting-state and structural connectivity findings in healthy samples of differential anterior-dorsal and posterior-dorsal precuneus connectivity (29,30). Specifically, stronger posterior hippocampal connectivity in PTSD for distal threats was localized to the posterior-dorsal precuneus subregion. By contrast, precuneal connectivity in MDD spanned the full anterior-posterior axis – weaker PAG– precuneus connectivity for near threats and stronger amygdala–precuneus connectivity for far threats. This finding is supported by Faul et al. (2020) using the same spatial fear conditioning paradigm in a healthy sample and found significant functional connectivity between the amygdala and anterior-dorsal precuneus during distal threat (5). Furthermore, the anterior-dorsal precuneus displayed increased activation to distal stimuli and reduced activation for the proximal CS+ stimulus in MDD. The posterior-dorsal precuneus exhibited sex-specific effects, displaying increased activity in higher MDD severity to proximal threats among females and decreased activity among males. This sex-divergent neural response may be due to the presentation of male avatars in the task.

These findings map onto precuneus subregion functions. The dorsal precuneus is a part of the parietal memory network and encodes complex visuospatial information (29,30). The posterior-dorsal precuneus is strongly connected to the early visual cortex and medial temporal memory systems to support visuospatial memory encoding, while the anterior-dorsal precuneus is preferentially connected to premotor and motor cortices and serves visuomotor spatial functions (30,31). In PTSD, our results suggest enhanced consolidation of threat-related visuospatial information between the posterior hippocampus and precuneus. MDD pathophysiology is driven by a broader disruption of both spatial mnemonic and sensorimotor integration that systematically varies by threat proximity. Stronger integration in the amygdala for distal threat may represent enhanced threat evaluation. Degraded integration within the PAG– precuneus circuit for proximal threats may reflect impaired information integration for mobilization of defensive action.

### Limitations and Strengths

The present study has several limitations. Our cohort was restricted to military veterans with predominantly combat-related index traumas. Consequently, the generalizability of these findings to civilian populations or distinct trauma types (e.g., interpersonal violence, chronic developmental trauma) remains to be established. The association with MDD symptoms severity may apply only to trauma-exposed individuals. Trauma exposure is highly prevalent in MDD (50-80%), but there is a significant subgroup of MDD patients who have not experienced trauma (32). While psychotropic medication use represents a potential confound, its impact is mitigated by the recruitment cap of 25% of our sample. Despite these limitations, the study possesses several notable strengths. Utilizing a 3D virtual reality paradigm allowed for the presentation of social threats in a naturalistic framework that helped enhance neural responses (33). An immersive environment enhances the ecological validity of our paradigm. Examining threats encountered at different spatial scales enhances our understanding of how threat networks are impacted post-trauma. Examining the temporal features of the fear learning process in relevant circuits will provide important information for intervening optimally in the peri-traumatic period.

## Conclusion

The human brain dynamically calibrates its responses to threats based on an egocentric spatiotemporal distance to threat. Distant threats engage the cognitive-fear network for strategic, evaluative processing, while proximal threats engage the reactive-fear network to mount immediate defensive responses. This dual architecture is disrupted among individuals with severe PTSD or MDD symptoms. Amygdala hyperreactivity emerged as uniquely associated with PTSD symptoms, predominately driven by hyperarousal symptoms, in response to distal threat. MDD symptom severity is associated with widespread alterations among the cognitive-fear network components in response to near threats. The observed pattern of brain activity suggests that severe post-trauma symptoms precipitate heightened safety signaling and altered spatial threat processing.

## Acknowledgements

This work was supported by a VA Merit Grant (*Brain Systems for Fear Generalization and Threat Processing in PTSD*) and a National Institute of Health R01 grant (MH 139648). We extend thanks to the Mid-Atlantic MIRECC for Post-Deployment Mental Health and to Leonard Faul for their support on this work.

