## Supplemental Data for "Dissociable Neural Responses to Conditioned Social Threats are Modulated by Spatial Proximity in PTSD and MDD"

**Supplementary Material**

**Supplementary Methods**

***MRI Data Acquisition***

Brain MRI data were acquired at the Duke-UNC Brain Imaging and Analysis Center. A significant hardware upgrade to the MRI scanner during data collection resulted in two sets of acquisition parameters and higher spatial resolution functional images for participants scanned after the upgrade (*n* = 25). Prior to the scanner upgrade, high-resolution 3D T1-weighted structural images were acquired using an axial Fast Spoiled Gradient-Recalled Echo (FSPGR) BRAVO sequence on a 3T GE SIGNA scanner equipped with an 8-channel head coil. The sequence parameters were as follows: repetition time (TR) = 8.16 ms; echo time (TE) = 3.18 ms; inversion time (TI) = 400 ms; flip angle = 12°; field of view (FOV) = 256 × 256 mm; acquisition matrix = 256 × 256; and 172 contiguous axial slices with a slice thickness of 1.0 mm. This yielded an isotropic voxel resolution of 1.0 × 1.0 × 1.0 mm³. Parallel imaging was employed with an acceleration factor of 2. The total receiver bandwidth was 62.5 kHz (244.1 Hz/voxel). Following installation of a new scanner, the T1-weighted images were acquired using an axial Magnetization-Prepared Rapid Gradient-Echo (MP-RAGE) sequence on a 3T GE SIGNA Premier scanner equipped with a 48-channel head and neck coil. The sequence parameters were: TR = 2379 ms; TE = 0.30 ms; TI = 750 ms; flip angle = 12°; acquisition matrix = 256 × 256; and contiguous axial slices with a slice thickness of 1.0 mm. Parallel imaging was utilized with an in-plane acceleration factor of 2. The voxel bandwidth was 244.1 Hz/voxel.

Prior to scanner upgrades, functional blood-oxygen-level-dependent (BOLD) images were acquired using an oblique axial SENSE-spiral sequence. The acquisition parameters were: TR = 2000 ms; TE = 27 ms; flip angle = 60°; FOV = 256 × 256 mm; and a reconstructed matrix of 64 × 64. Thirty-four interleaved slices were acquired with a slice thickness of 3.8 mm, yielding an effective voxel resolution of 4.0 × 4.0 × 3.8 mm³. Following scanner upgrade, functional BOLD images were acquired using a 2D gradient-echo echo-planar imaging (EPI) sequence. The acquisition parameters were: TR = 2000 ms; TE = 25 ms; flip angle = 52°; acquisition matrix = 128 × 128; and 69 contiguous slices with a slice thickness of 2.0 mm, yielding an effective voxel resolution of 2.0 × 2.0 × 2.0 mm³. Simultaneous multi-slice (GE HyperBand) acquisition was employed with an acceleration factor of 3. In-plane parallel imaging was concurrently applied with an acceleration factor of 2. Phase encoding was performed in the anterior-to-posterior (AP) direction. The sequence utilized an effective echo spacing of 0.30 ms and a total readout time of 37.8 ms.

***MRI Data Preprocessing***

Anatomical T1-weighted MRI data was processed with FreeSurfer’s recon-all (1). White matter and cerebrospinal fluid (CSF) segmentation masks were used to extract signals for physiological signal regression (aCompCor) (2). Functional BOLD data was preprocessed using FSL 6.0.7 (FMRIB Software Library, www.fmrib.ox.ac.uk/fsl). Preprocessing steps included field map correction, removal of the first 4 volumes to allow for magnetization equilibrium, motion correction using MCFLIRT (3), grand-mean intensity normalization, highpass temporal filtering, and spatial smoothing (5.0 mm FWHM). Registration to high resolution structural and standard space (MNI152NLin6Asym) images was carried out using FLIRT. Global smoothness of brain data was estimated using AFNI’s 3dFWHMx (4).

***Univariate Analysis***

First-level general linear models (GLMs) were implemented in FSL's FEAT. Task regressors were convolved with the double-gamma hemodynamic response function and entered alongside 24 motion parameters (6 realignment parameters, their temporal derivatives, and squared terms) and 10 aCompCor components (5 white matter, 5 CSF) as nuisance regressors. Univariate hemodynamic response associations with PTSD and MDD symptom severity were examined within threat processing regions using linear mixed-effects models (see *Statistical Analysis*).

***Statistical Thresholding and Multiple Comparison Correction***

Cluster-based thresholding was performed using AFNI’s 3dClustSim and 3dClusterize (5), with clusters defined as contiguous voxels sharing at least one face or edge. Cluster-extent thresholds were determined using a two-tailed voxelwise threshold of *p* < 0.001 (uncorrected) and a familywise error (FWE) rate of α = 0.05 for univariate analyses. For gPPI analyses, the α threshold was Bonferroni-corrected for the five seeds to 0.05 / 5 . Given the small volumes of subcortical structures of interest, a separate small-volume correction (SVC) was applied to the bilateral amygdala, hippocampus, thalamus, and PAG. A subcortical mask was used to estimate the cluster-extent threshold required to maintain *p_FWE_* < 0.05 within the subcortical regions, while retaining the global smoothness estimate from the broader region-of-interest mask to preserve the integrity of spatial autocorrelation modeling.

***Dominance Analysis***

Dominance analysis was conducted to determine the relative importance of regressors using the *domir* package in R (6). This analysis was performed to better understand the unique contributions of individual PTSD and MDD symptoms on neural responses to threat. The average contrast of parameter estimates (CS+ > CS-) from each significant univariate and gPPI voxel cluster served as the dependent variables. PTSD models evaluated the relative importance of the four CAPS-5 symptom domains (re-experiencing, avoidance, negative alterations in mood/cognition, and hyperarousal), while MDD models evaluated the cognitive and somato-affective dimensions of the BDI-II. Because dominance analysis iteratively compares all possible combinations of predictors, age and sex were maintained as fixed covariates across every resulting submodel to isolate the effects of the clinical variables. We assessed three levels of dominance with increasing stringency: general, conditional, and complete dominance. Symptoms were ranked based on their overall R^2^ contribution (general dominance), their average incremental R^2^ contribution within models of a specific size (conditional dominance), and whether their incremental contribution strictly exceeded that of another predictor across every possible submodel (complete dominance).

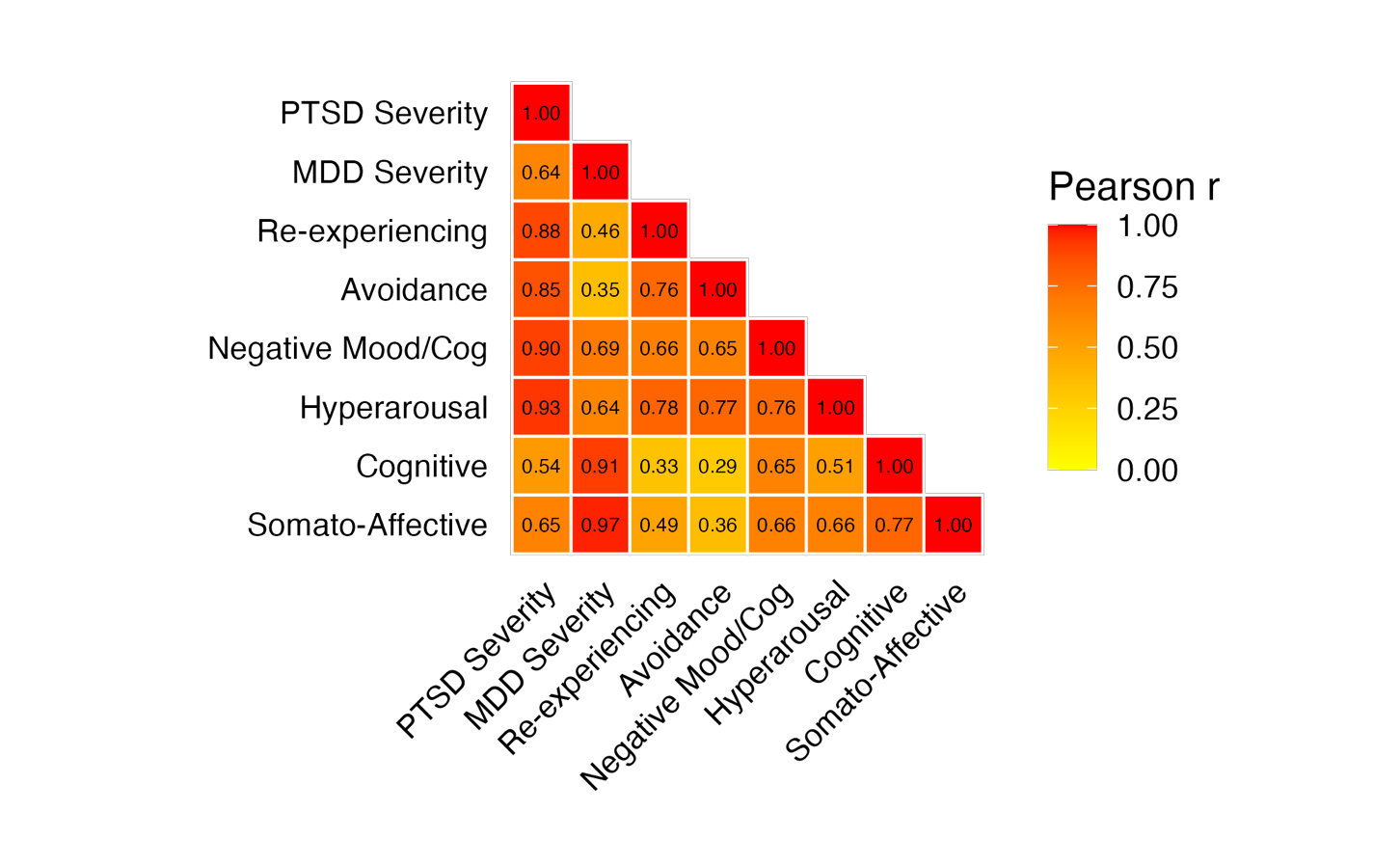

**Figure S1**: Clinical variable correlation plot.

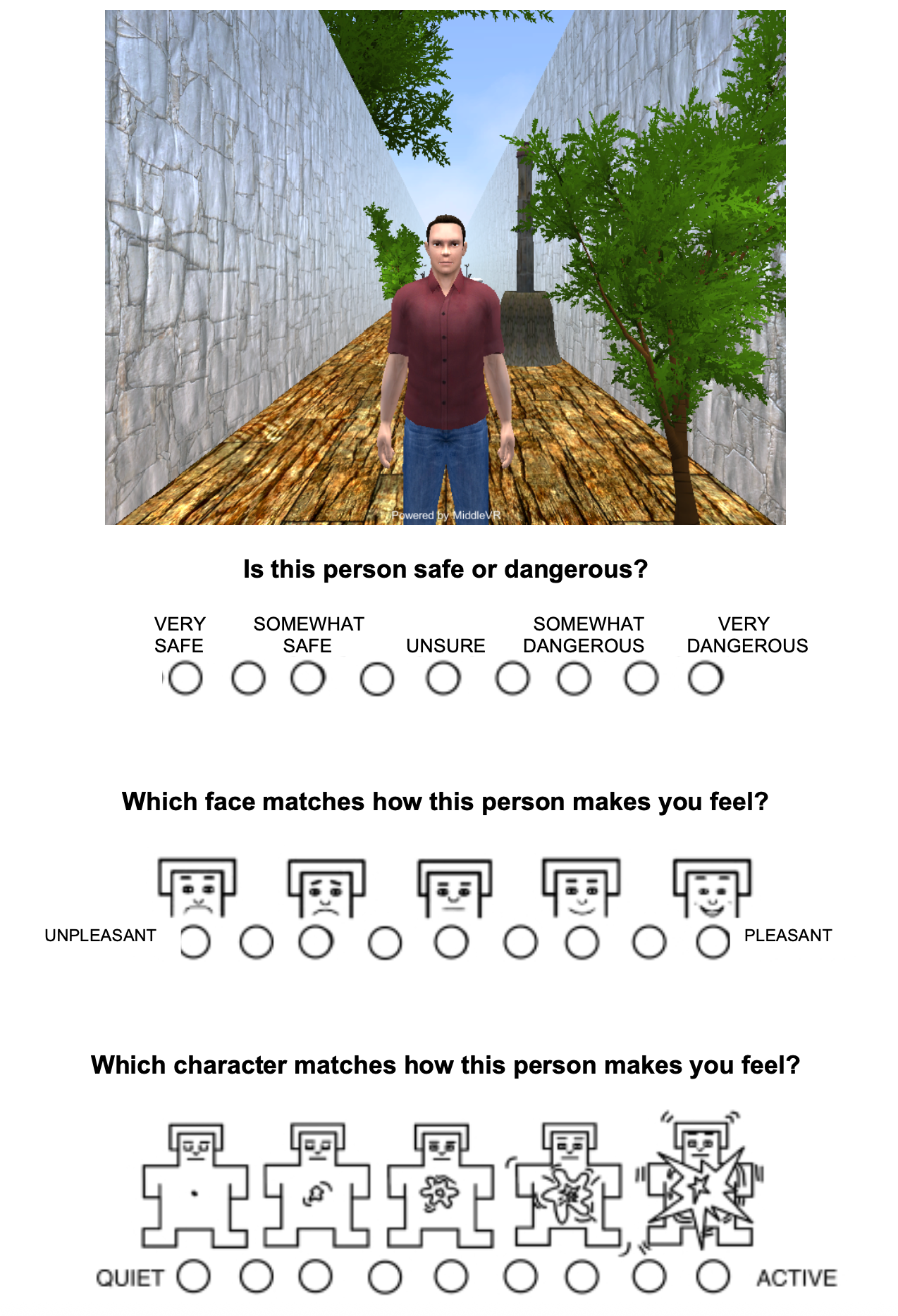

**Figure S2.** Example post-scan questionnaire

**Table S1.** Arousal rating comparisons

| **Rating 1** | **Rating 2** | **df** | **t-statistic** | **p** |
| --- | --- | --- | --- | --- |
| CS-_Far_ | CS-_Near_ | 46 | 0.64 | 5.26E-01 |
| CS-_Far_ | CS+_Far_ | 46 | -5.51 | 1.54E-06 |
| CS-_Far_ | CS+_Near_ | 46 | -9.27 | 4.22E-12 |
| CS-_Near_ | CS+_Far_ | 46 | -6.08 | 2.22E-07 |
| CS-_Near_ | CS+_Near_ | 46 | -8.62 | 3.67E-11 |
| CS+_Far_ | CS+_Near_ | 46 | -5.54 | 1.42E-06 |

**Table S2.** Valence rating comparisons

| **Rating 1** | **Rating 2** | **df** | **t-statistic** | **p** |
| --- | --- | --- | --- | --- |
| CS-_Far_ | CS-_Near_ | 46 | -0.21 | 8.31E-01 |
| CS-_Far_ | CS+_Far_ | 46 | 6.03 | 2.65E-07 |
| CS-_Far_ | CS+_Near_ | 46 | 8.47 | 6.01E-11 |
| CS-_Near_ | CS+_Far_ | 46 | 5.34 | 2.80E-06 |
| CS-_Near_ | CS+_Near_ | 46 | 7.88 | 4.41E-10 |
| CS+_Far_ | CS+_Near_ | 46 | 4.86 | 1.39E-05 |

**Table S3.** Linear mixed model results including both PTSD and MDD symptom severity in the same model.

| **Region** | **Distance** | **Time** | **Volume (mm³)** | **Mean Z** | **Max Z** | **Peak MNI (x, y, z)** |
| --- | --- | --- | --- | --- | --- | --- |
| ***PTSD Severity Main Effect*** | | | | | | |
| Ventral Posterolateral Thalamus L.* | Near | Late Acquisition | 200 | -3.75 | -4.43 | -16, -22, 0 |
| Precentral Gyrus L. | Far | Late Acquisition | 320 | -3.65 | -4.35 | -26, -14, 64 |
| ***MDD Severity Main Effect*** | | | | | | |
| Hippocampus L.* | Far | Early Acquisition | 128 | 4.28 | 5.29 | -28, -30, -10 |
| Dorsolateral Prefrontal Cortex R. | Near | Late Acquisition | 672 | -3.93 | -5.61 | 38, 26, 42 |
| Supplementary Motor Area | Far | Late Acquisition | 744 | 4.10 | 5.42 | 0, -10, 50 |
| Precentral Gyrus L. | Far | Late Acquisition | 392 | 3.62 | 4.24 | -26, -14, 64 |

*Notes: ** *Regions were small volume corrected (SVC)*

**PTSD Diagnosis Results**

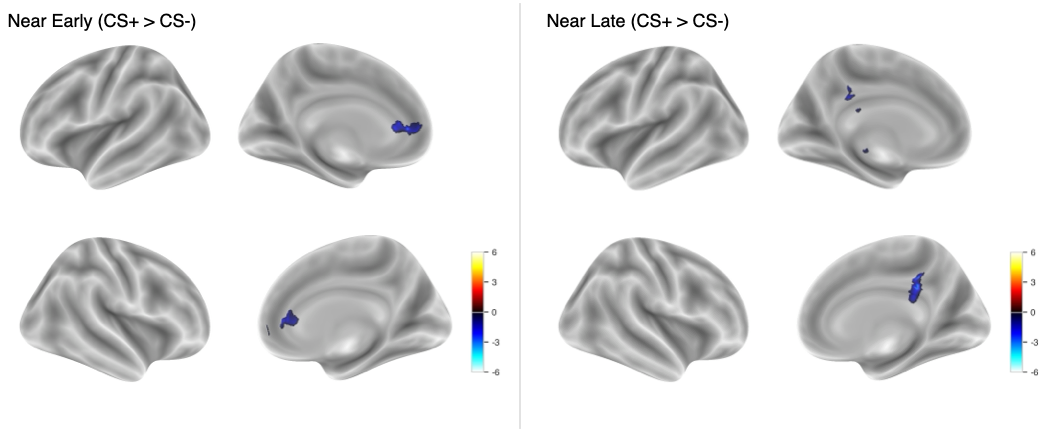

**Figure S3**: Associations between PTSD diagnosis and neural activation for the CS+ > CS- contrasts.

**MDD Diagnosis Results**

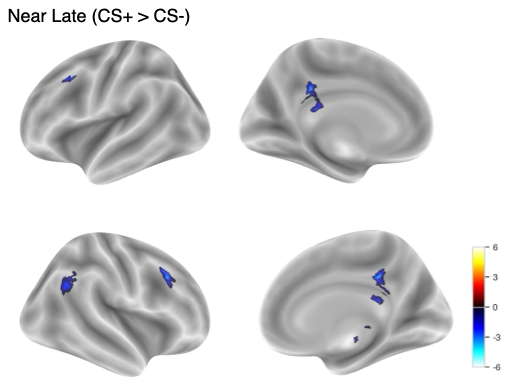

**Figure S4**: Associations between MDD diagnosis and neural activation for the near threat CS+ > CS- contrast during late acquisition.
